# α2δ-2 protein controls structure and function at the cerebellar climbing fiber synapse

**DOI:** 10.1101/685818

**Authors:** Kathleen A. Beeson, Ryne Beeson, Gary L. Westbrook, Eric Schnell

## Abstract

α2δ proteins (*Cacna2d1-4*) are auxiliary subunits of voltage-dependent calcium channels that also drive synapse formation and maturation. Because cerebellar Purkinje cells (PCs) only express one isoform of this family, α2δ-2 (*Cacna2d2*), we used PCs as a model system to examine roles of α2δ in excitatory synaptic function in a *Cacna2d2* knockout mouse. Whole-cell recordings of PCs from acute cerebellar slices revealed altered climbing fiber (CF)-evoked complex spike generation, as well as increased amplitude and faster decay of CF-evoked excitatory postsynaptic currents (EPSCs). CF terminals in the KO were localized more proximally on PC dendrites, as indicated by VGLUT2^+^ immunoreactive puncta, and computational modeling demonstrated that the increased EPSC amplitude can be partly attributed to the more proximal location of CF terminals. In addition, CFs in KO mice exhibited increased multivesicular transmission, corresponding to greater sustained responses during repetitive stimulation, despite a reduction in the measured probability of release. Electron microscopy demonstrated that mutant CF terminals had twice as many vesicle release sites, providing a morphologic explanation for the enhanced glutamate release. Though KO CFs evoked larger amplitude EPSCs, the charge transfer was the same as wildtype as a result of increased glutamate re-uptake, producing faster decay kinetics. Together, the larger, faster EPSCs in the KO explain the altered complex spike responses, which degrade information transfer from PCs and likely contribute to ataxia in *Cacna2d2* KO mice. Our results also illustrate the multidimensional synaptic roles of α2δ proteins.

**Significance Statement:** α2δ proteins (*Cacna2d1-4*) regulate synaptic transmission and synaptogenesis, but co-expression of multiple α2δ isoforms has obscured a clear understanding of how various α2δ proteins control synaptic function. We focused on roles of the α2δ-2 protein (*Cacna2d2*), whose deletion causes cerebellar ataxia and epilepsy in mice and humans. Because cerebellar Purkinje cells only expresses this single isoform, we studied excitatory climbing fiber synaptic function onto Purkinje cells in *Cacna2d2* knockout mice. Using optical and electrophysiological analysis, we provide a detailed description of the changes in Purkinje cells lacking α2δ-2, and provide a comprehensive mechanistic explanation for how functional synaptic phenotypes contribute to the altered cerebellar output.

## Introduction

Synapses are indispensable to neural circuit function, yet our understanding of synapse formation and physiology in health and neurological disease is incomplete. Recently, α2δ proteins (*Cacna2d1-4*) have been recognized as important regulators of synapse formation and plasticity (Dolphin, 2012). In addition to their roles in synaptic transmission as auxiliary subunits of voltage-dependent calcium channels (VDCCs) (Canti et al., 2005; Hoppa et al., 2012), these proteins have multiple postsynaptic roles in driving synapse formation, trans-synaptic communication, and glutamate receptor function (Eroglu et al., 2009; Kurshan et al., 2009; Fell et al., 2016; Wang et al., 2016; Brockhaus et al., 2018; Chen et al., 2018; Geisler et al., 2019). Mutations in human α2δ genes have been associated with epilepsy, movement disorders, and schizophrenia (Edvardson et al., 2013; Pippucci et al., 2013), and α2δ-1/2 proteins are the primary targets of the widely prescribed antiepileptic and analgesic, gabapentin (Gee et al., 1996; Brown and Gee, 1998; Boning Gao, 2000). However, the roles of α2δ proteins in controlling synaptic and network function at any given synapse are still elusive, due in part to expression of multiple isoforms in many neurons.

Purkinje cells (PCs), the primary output pathway from the cerebellar cortex, exclusively and abundantly express the α2δ-2 isoform (*Cacna2d2*) (Barclay et al., 2001a; Cole et al., 2005; Lein et al., 2007; Dolphin, 2012), thus providing an opportunity to determine how this protein contributes to synaptic transmission. Like the spontaneous ‘ducky’ mouse mutants (*du^2j^/du^2j^* or *du/du*), targeted deletion of α2δ-2 causes cerebellar ataxia, epilepsy and premature death (Barclay et al., 2001b; Brodbeck et al., 2002; Ivanov et al., 2004; Donato et al., 2006), indicative of its importance in neurological function.

The global loss of α2δ-2 likely impacts synapses throughout the brain. However, the climbing fiber (CF) to PC synapse has multiple distinctive features, including the mono-innervation of mature PCs by a single CF (Hashimoto et al., 2009) and presynaptic expression of vesicular glutamate transporter 2 (VGLUT2), allowing for the visualization of CF terminals and making this an ideal site to determine how α2δ-2 contributes to synaptic function. In contrast to Purkinje cells, the inferior olivary cells (from which CFs arise) predominantly express the α2δ-1 isoform, with low expression of α2δ-2/3 isoforms (Cole et al., 2005; Lein et al., 2007). Finally, CF activity drives complex spike (CpS) generation in PCs, which produces a high-fidelity error prediction signal important to the processing of motor coordination and learning (Yang and Lisberger, 2014; Heffley et al., 2018).

We combined structural and electrophysiological analysis of the CF to PC synapse in *Cacna2d2* knockout mice (Ivanov et al., 2004) to elucidate the contribution of α2δ-2 to excitatory synapse formation and transmission. Contrary to positive regulation of excitatory synaptogenesis by the α2δ-1 isoform (Li et al., 2004; Eroglu et al., 2009; Chen et al., 2018; Risher et al., 2018), loss of α2δ-2 increased functional synaptic innervation by CFs. α2δ-2 KO CFs had elevated glutamate release and clearance compared to WT, resulting in profound deficiencies in the generation of CpS spikelets. Together, our studies demonstrate the critical role of α2δ-2 in proper CF-PC synapse organization and network function, and allude to the wide versatility of α2δ proteins in synaptic transmission.

## Materials & Methods

### Animals

*Cacna2d2* knockout mice (*Cacna2d2^tm1Svi^*, MGI = 3055290; generously supplied by Drs. Sergey Ivanov and Lino Tessarollo) were obtained as cryopreserved sperm and re-derived via *in vitro* fertilization on a C57B/6J background. Breeding mice were kept heterozygous, and genotyping was performed using the following primers: forward 5’-ACTGGTGGGCATCTTACAGC-3’, reverse mutant 5’ - AAAGAACGGAGCCGGTTG-3’, reverse wildtype 5’-TGTTAGCGGGGAGGTCACTA-3’. This produced a ∼700 bp product from the wildtype allele and a ∼550 bp product from the mutant allele. Mice were born in the Mendelian ratio of 1:2:1; with WT mice having two copies of the intact *Cacna2d2* gene, and KO mice as homozygous mutants. Because ∼50% of KO mice die prematurely (Ivanov et al., 2004), all experiments used male and female mice between p21-p30, when CF-PC synapses have reached maturity, but before significant loss of KO mice.

*Cacna2d2* deletion was validated, and *Cacna2d* isoform abundance were measured, using quantitative PCR from fresh-frozen tissues. Regions of the inferior olive (IO) and the Purkinje cell layer (PCL), including the proximal molecular layer, were microdissected by laser capture for pre- vs. postsynaptic comparative analysis. Briefly, mice were deeply anesthetized by inhalation of 4% isoflurane followed by injection of 0.8 ml of 2% avertin i.p. (Sigma-Aldrich) and rapidly decapitated. Brains were rapidly dissected, mounted in OCT medium (Tissue-Tek) on dry ice, and placed at −20°C for cryosectioning. 25 µm coronal sections from cerebellum and brainstem were collected on RNase-treated PEN membrane slides (Zeiss). Slides were then dehydrated through a succession of EtOH rinses (70% EtOH 30 seconds, 100% EtOH 30 seconds), and nuclei stained using 1 mg/mL Cresyl Violet in 100% EtOH (1 minute). IO and PCL regions were identified based on anatomic criteria, and isolated using laser microdissection (Zeiss). Samples were resuspended in 20 µl QIAzol Lysis Reagent (Qiagen) for 20 minutes at RT and stored at −80°C. RNA isolation and qPCR were performed by the Oregon Health & Science University Gene Profiling/RNA and DNA Services Shared Resource. In brief, RNA was isolated using the Trizol/RNeasy hybrid protocol with QIAcube automation. SuperScript™ IV VILO™ Master Mix (Invitrogen) was used for reverse transcription with 636 pg of input RNA per 20 µl reaction. Following reverse transcription, cDNA was quantified and normalized for 7500ng of cDNA input for pre-amplification in 14 PCR cycles. A 1:4 dilution of the pre-amp reaction was used as input for qPCR. The TaqMan Universal master mix (Life Technologies) was used for the PCR reaction, using a single master mix per TaqMan probe set for *Cacna2d1* (Mm00486607_m1), *Cacna2d2* (Mm01230564_g1), *Cacna2d3* (Mm00486641_m1), *Cacna2d4* (Mm01190080_m1), and *ACTB* (Mm04394036_g1) as the endogenous control. Data were acquired using Applied Biosystems QuantStudio 12K Flex Software v1.2.2 (Life Technologies) with settings set to default. Measurements are reported as either ΔCt values (difference in cycle time for gene of interest relative to ACTB, in which a higher ΔCt indicates lower relative expression) or mean fold difference, as determined by 2^^-(Ct target – Ct ACTB target) – (Ct reference – Ct ACTB reference)^ ± SEM.

Mice were maintained in facilities fully accredited by the Association for Assessment and Accreditation of Laboratory Animal Care and veterinary care was provided by Oregon Health & Science University’s Department of Comparative Medicine. All animal care and experiments were performed in accordance with state and federal guidelines, and all protocols were approved by the OHSU Institutional Animal Care and Use Committee.

### Immunohistochemistry

Following deep anesthesia as described above, p21 *Cacna2d2* KO and WT littermates were transcardially perfused with 5 ml phosphate-buffered saline (PBS) followed by 4% paraformaldehyde-PBS (PFA-PBS). Mice were decapitated, and brains were removed and post-fixed for 24 hrs. in 4% PFA-PBS. Cerebella were cut in sagittal sections at 50 µm on a vibratome and stored in PBS at 4°C. Sections containing vermis lobe VI were permeabilized with 0.4% Triton-PBS containing 10% normal goat serum for 1hr at room temperature, then stained with mouse anti-Calbindin 1:20 (Neuromab #73-452) and guinea pig anti-VGLUT2 1:200 (Synaptic Systems #135404) at 4°C overnight. After rinsing, corresponding fluorescently-labeled secondaries (Invitrogen) were applied at 1:400 and glass coverslips were mounted on glass slides with Fluoromount G (Sigma-Aldrich).

To image VGLUT2^+^ synapses, 6 µm z-stack images from the vermis lobe VI were acquired at 0.2 µm intervals with a Zeiss LSM780 laser scanning confocal microscope at 40x magnification and 1024 x 1024 pixel density using ZEN software. This produced images of the entire thickness of the molecular layer, including the PC somata. Images were then analyzed in FIJI/ImageJ. VGLUT2^+^ puncta distribution/density were quantified from maximum projection images as distinct VGLUT2^+^ puncta of at least 0.2 µm^2^, after subtracting the background for increased contrast. For each punctum, the y-distance from the top of the PC layer to the terminal was measured. Per image, a minimum of 100 µm of linear PC layer was quantified. VGLUT2^+^ puncta distributions were then binned into 10 µm distances. VGLUT2^+^ puncta size and a second measure of density (from whole volume of tissue) were calculated using a masking feature in FIJI/ImageJ that captures puncta between 0.1-5µm^2^, and this was normalized to the length of PCL imaged to produce a measure of density per µm of PCL. Punctum distribution, density and size data were averaged from 2-3 sections/animal, using 5-6 animals of each genotype. Slides were coded prior to imaging, and image acquisition and analysis were performed by investigators blinded to genotype. Sections were stained side-by-side with the same antibody mixtures, and imaging parameters were kept constant between samples.

### Slice Preparation and Electrophysiology

KO or WT littermates were deeply anesthetized as described above, and transcardially perfused with ice-cold sucrose-based solution containing (mM): 87 NaCl, 2.5 KCl, 1.25NaH_2_PO_4_, 0.4 ascorbate, 2 Na pyruvate, 25 D-glucose, 25 NaHCO_3_, 75 Sucrose, 7 MgCl_2_, 0.5 CaCl_2_ (osmolarity adjusted to 300-305 mOsm) and equilibrated with 95% O_2_ and 5% CO_2_ gas mixture. Acute 300 µm sagittal slices were cut from cerebellum using a vibratome (VT1200, Leica Microsystems), and incubated for 30 minutes in standard artificial cerebral spinal fluid (aCSF) at 34°C.

Whole-cell recordings were obtained using 1-3 MΩ borosilicate glass pipettes filled with either K-gluconate or CsCl_2_-based internal solution. For current clamp experiments, internal solution contained (in mM): 135 K-gluconate, 10 HEPES, 10 NaCl, 1 MgCl_2_, 0.1 BAPTA, 0.1 CaCl_2_, 2 ATP-Mg, and 10 phosphocreatine, pH 7.28 adjusted with KOH (osmolarity adjusted to 289 mOsm). For voltage clamp recordings, the internal solution contained (in mM): 100 CsMeSO_4_, 35 CsCl, 15 TEA-Cl, 1 MgCl_2_, 15 HEPES, 0.2 EGTA, 2 ATP-Mg, 0.3 TrisGTP, 10 phosphocreatine, and 2 QX-314, pH 7.3 adjusted with CsOH (osmolarity adjusted to 295 mOsm). External solution contained (in mM): 125 NaCl, 25 NaHCO_3_, 1.25 NaH_2_PO_4_, 3 KCl, 25 Dextrose, 2 CaCl_2_, 1 MgCl_2_ (osmolarity adjusted to 300 mOsm) and was continuously perfused via roller pump.

PCs from the vermis, lobe VI, were identified by soma size and location in the PC layer using live infrared differential contrast microscopy on an upright Olympus microscope. Inhibition was blocked in all experiments by 10 µM SR95531 (Tocris). Whole-cell patch-clamp recordings were obtained in voltage clamp mode with cell capacitance, series resistance and input resistance monitored in real time using intermittent −10 mV voltage steps. Signals were amplified with a MultiClamp 700B (Molecular Devices) amplifier and pipette capacitance was compensated for using MultiClamp software. Signals were low-pass filtered at 6 kHz and sampled at 10 kHz, and digitized with a National Instruments analog-to-digital board. All recordings were acquired and analyzed using IgorPro-based (Wavemetrics) software. All recordings were performed at room temperature.

Climbing fiber-mediated excitatory postsynaptic currents (EPSCs) were evoked using theta or monopolar glass electrode stimulation in the granule cell layer (0.1 ms, 0-100 V square pulses; 0.1 or 0.05 Hz stimulation frequency), placed ∼50 µm from the PC, and the stimulation electrode position was adjusted as needed to obtain a CF response. CF responses were identified as large all-or-none EPSCs that appeared during incremental increases in stimulation intensity, with paired-pulse depression when stimulated with 2 pulses 50 ms apart. All cells were first recorded in voltage clamp mode, where a hyperpolarizing step of −10 mV was applied to monitor cell capacitance, series resistance and input resistance. Series resistance was not compensated; cells with series resistance >10 MΩ, or a >2 MΩ change in series resistance over the course of the experiment, were excluded.

For current clamp experiments, PC resting membrane potential was measured by holding the cell in zero current mode, then a small amount of bias current (approximately −130 pA) was injected to keep the cell near −60 mV for complex spike and current step experiments. Bridge balance was applied to compensate for pipette and series resistance throughout the recording. First, CFs were stimulated at 0.1 Hz and 10 consecutive sweeps of evoked complex spikes were recorded. Then with CF stimulation off, 500 ms steps of current injection from −200 pA– 1 nA were delivered at least 5 times per step. Cells were returned to voltage clamp mode to assess recording stability. For complex spike analysis, sweeps were averaged to measure the initial spike amplitude and rise rate. Spikelets were classified as rapid depolarizations (>1000 V/s) that reached +20 mV from baseline, from which spikelet trough-to-peak amplitudes were measured. Area under the curve was measured by integrating the trace during the first 100 ms following stimulation.

For most voltage clamp experiments, cells were held at −70 mV in the presence of 0.5 µM NBQX (Tocris) to maintain voltage clamp during CF EPSCs. Brief hyperpolarizing steps (−10 mV) were delivered to monitor PC access and input resistance preceding each CF-evoked EPSC. 10 CF EPSC trials were averaged, and these averages were used to determine peak amplitude, 20-80% rise time and tau of decay (from single-exponential fits of the EPSC decay). Then, ten paired-pulse responses with an inter-stimulus interval of 50 ms were collected, followed by 10 Hz trains of CF stimulation or drug wash-in experiments. For DL-TBOA experiments, baseline EPSCs were acquired at 0.05 Hz before 50 µM DL-TBOA (Tocris) was added to the bath. For kynurenic acid (KYN) experiments, aCSF excluded NBQX and PCs were held at −20 mV to maintain voltage-clamp during 0.1 Hz CF stimulation. After acquiring baseline EPSCs, 1 mM KYN (Sigma-Aldrich) was added to the perfusate. Quantification of EPSC peak amplitude and tau of decay from drug wash-in experiments used averages from 10 sweeps prior to wash-on compared to 10 sweeps after 10 minutes of exposure to drug (For KYN effect: (EPSC_Control_ - EPSC_KYN_) * 100; For TBOA effect: (τ_TBOA_ - τ_Control_) * 100). For asynchronous EPSC (aEPSC) experiments, aCSF was composed of 1.3 mM Sr^2+^ in replacement of Ca^2+^, 3.3 mM Mg^2+^, and NBQX was omitted. PCs were held at −70 mV and CFs were stimulated at 0.05 Hz. ∼10 trials were used for quantification of aEPSCs, which were sampled from a 500 ms window starting at 150 ms from CF stimulation, and selected manually. For data presentation, aEPSC traces were off-line box-filtered at 1 kHz. To estimate the readily releasable pool, cumulative CF response amplitudes were plotted, and the last third of the train was fit with a linear regression that was extrapolated to time 0 (Schneggenburger et al., 1999).

### NEURON Computational PC Model

CF simulations were performed with NEURON version 7.7.0, using source code generously supplied by Dr. Michael Häusser (Roth and Häusser, 2001). The following model parameters were kept constant across all simulations: R_m_ = 120.2 kΩcm^2^, C_m_ = 0.64 µF/cm^2^ and a residual uncompensated series resistance of 1 MΩ. Because the Model is based on recordings and dimensions of a p21 rat PC (Roth and Häusser, 2001), we normalized our WT measurements for CF distribution to the model cell as control, and adjusted the dendritic innervation by KO CFs based on our VGLUT2^+^ distribution as a relative decrease in length of coverage (i.e. 0.7x CF length of control). CF EPSCs were simulated using 500 inputs of 1 nS peak conductance (simulated as the sum of two exponentials for rise and decay) with a reversal potential of 0 mV and a constant density per dendritic length distribution. Simulation time step was 10 µs for integration. Waveforms were created in IgorPro8 from simulation output to measure EPSC decay time constants by fitting with a single exponential.

### Transmission Electron Microscopy (TEM)

Animals were deeply anesthetized with isoflurane and avertin, as described above, and then transcardially perfused with 10 ml ice-cold heparin (1k usp per ml; Novaplus) followed by freshly prepared 2% glutaraldehyde/2% paraformaldehyde in 0.1M PB solution filtered with #3 filter paper (VWR) and pH to 7.4. Brains were dissected, cerebella were blocked, and post-fixed for 30 minutes in 4% paraformaldehyde. Tissue was transferred to 0.1 M PB for storage at 4°C and 40 µm sagittal slices of vermis lobe VI were made using a Leica microtome. Slices were collected in separate wells to assure TEM would be from slices >100 µm apart. PFA, Glutaraldehyde and microtome blades were all from Electron Microscopy Sciences (EMS).

Tissue samples were coded before processing for TEM by a blinded investigator. Briefly, sections were incubated in 1% osmium tetroxide in PB for 30 minutes, dehydrated through a graded series of ethanols, placed into propylene oxide for 30 min, and then placed in 1:1 propylene oxide:EMBed resin (EMS) rotating at room temperature overnight. Sections were then incubated in 100% resin for 2 hrs, embedded between sheets of Aclar plastic, and incubated at 60°C for 48 hrs. Cerebellum sections were then glued to resin blocks and ultrathin sections (50 nm) were collected onto 400 mesh copper grids (EMS). The ultrathin sections were then counterstained with uranyl acetate and Reynolds lead citrate and examined using a FEI Tecnai 12 electron microscope (Hillsboro, OR) and images were captured using an Advanced Microscopy Techniques digital camera.

CF terminals were imaged from the most highly innervated region of the molecular layer (20-60 µm from PC somata) and identified at 6800x magnification by the following criteria: proximity (<3 µm) to PC primary dendrites, dense-packing of round vesicles, and when synaptic contacts were present, asymmetric Gray’s type-1 excitatory synaptic markers. Images were taken at 18500x magnification and CF terminals with clearly delineated membranes that met the criteria described above were analyzed using Fiji/ImageJ software by a separate blinded investigator. Terminal area, total SV density, the length and number of synaptic contacts made by each terminal (as determined by the opposing postsynaptic density), as well as the number of SVs within 100 nm of each synaptic contact as a proxy for the readily releasable pool were measured. Quantifications from ∼10-15 CF images/animal were averaged (n = animal).

### Statistics

Data was tested for Gaussian distribution using the Kolmogorov-Smirnov normality test with Dallal-Wilkinson-Lilliefor P value. Differences between genotypes in VGLUT2^+^ puncta distribution and TEM total synaptic contacts/terminal were tested for significance using Kolmogorov-Smirnov test. For other nonparametric data, the Mann-Whitney test was used (i.e. CpS spikelet number). Immunofluorescence, TEM, and electrophysiology data with normal distributions were analyzed using Student’s t-tests. Normalized and cumulative amplitude responses during repetitive CF stimulation were compared between genotypes using Multiple t-tests with Holm-Sidak correction. For morphology data, multiple images per animal were averaged where n = # mice. For electrophysiology, all experiments utilized at least 4 animals per genotype, where n = # cells. Data were graphed in Prism GraphPad version 8 and are reported as the mean ± SEM. *p<0.05, **p<0.01, ***p<0.001, ****p<0.0001.

## Results

### α2δ-2 controls Purkinje cell spiking patterns in response to climbing fiber activation

To examine how the loss of α2δ-2 affects cerebellar output, we performed whole-cell recordings from PCs in acutely prepared cerebellar slices from α2δ-2 knockout (KO) mice and their wildtype (WT) littermates at p21-p30, after CF innervation has reached maturity (Hashimoto et al., 2009). We focally stimulated CFs in the granule cell layer and unitary CF-mediated EPSCs were identified by their large amplitude, all-or-nothing nature and paired-pulse depression (Dittman and Regehr, 1998; Hashimoto and Kano, 1998; Liu and Friel, 2008; Rudolph et al., 2011). CF-evoked CpSs were then recorded in current clamp mode.

The voltage envelope of CpS waveforms was comparable between WT and KO (Figure 1A; Integral of CpS between time 0 - 100 ms: WT = 0.96 ± 0.06 V • s, n = 10; KO = 0.81 ± 0.06 V • s, n = 11; p = 0.1; unpaired Student’s t-test), but the number of CpS spikelets was substantially reduced in KO PCs (Figure 1B; WT = 3.2 ± 0.5, n = 10; KO = 1.2 ± 0.4, n = 11; p = 0.002; Mann-Whitney). In addition, the few spikelets that did occur during the KO CpS were of lower trough-to-peak amplitude. All first spikelets in the WT were >30 mV compared to 75% in the KO (Figure 1C), which is consistent with a lower probability of CpS transmission (Khaliq and Raman, 2005) in the α2δ-2 KO. Moreover, though subsequent spikelets were present in 2 of 11 KO cells, none reached 30 mV trough-to-peak amplitude (Mean KO spikelet_2_ = 16.5 ± 5.7 mV, n = 2; spikelet_3_ = 7.6 mV, n = 1; spikelet_4_ = 7.2 mV, n = 1; spikelet_5_ = 6.5 mV, n = 1). Despite the differences in CpS spikelet generation between genotypes, initial spike amplitudes (Figure 1D) and rise rates were unchanged between WT and KO (Spike slope: WT = 1790 ± 230 V/s, n = 10; KO = 1660 ± 240 V/s, n = 11; p = 0.69; unpaired Student’s t-test).

**Figure 1.**
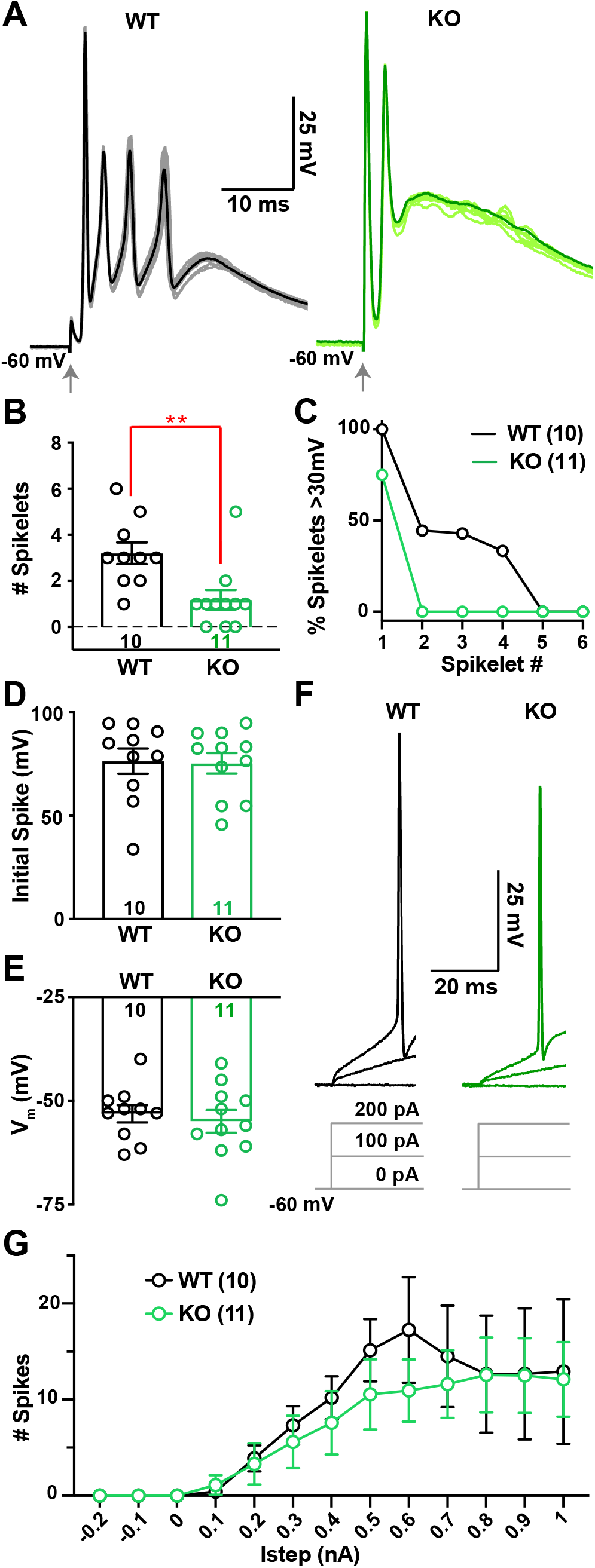
Climbing fiber (CF)-evoked complex spikes (CpSs) are altered in the Cacna2d2 KO, but intrinsic PC excitability is unchanged. (*A*) Representative CF-evoked complex spikes in WT (left) and KO (right) PCs, arrow indicates CF stimulation. Each trace consists of 10 overlaid traces (lighter color) and the corresponding CpS average (dark color). (*B*) Average number of spikelets per CpS in WT and KO PCs; **p < 0.01 (Mann-Whitney). (*C*) Percentage of spikelets exceeding 30 mV trough-to-peak amplitude, ordered by spikelet number. (*D*) Average CpS initial spike amplitude; p = 0.89 (NS). (*E*) Average PC membrane potential when in zero current mode; p = 0.60 (NS). (*F*) Representative single traces of membrane voltage responses to current injection, showing steps of 0, 100, 200 pA injections from V_m_ = −60 mV; *Left* WT (black); *Right* KO (green). Average I_step_ to initiate spiking WT = 200 ± 39 pA, n = 10; KO = 289 ± 82 pA, n = 9; p = 0.33 (NS). (*G*) Average spike count during current steps from −0.2 to 1 nA, V_m_ = −60 mV. WT (black) and KO (green); p = 0.63 (NS; Two-way ANOVA with repeated measures, F_(1, 19)_ = 0.23). Data shown ± SEM, n = cells; unpaired Student’s t-test unless otherwise indicated.

The altered CpS in KO PCs were not associated with changes in the intrinsic excitability of α2δ-2 KO PCs as measured by their resting membrane potentials (Figure 1E), input resistance (WT 140 ± 20 MΩ, n = 8; KO 122 ± 7.8 MΩ, n = 11; p = 0.38, unpaired Student’s t-test), or response to current injection (Figures 1F and 1G). Thus, we explored whether altered CpS patterns might represent differences in CF-mediated synaptic currents.

### α2δ-2 knockout mice have larger CF-evoked EPSCs with accelerated decay kinetics

The kinetics of CF-evoked excitatory postsynaptic currents (EPSCs) influence the CpS shape, such that a slower EPSC decay increases the likelihood of spikelet generation (Rudolph et al., 2011). Spikelet generation is also sensitive to the peak synaptic conductance, as an increased phasic conductance can result in depolarization block and failure to generate spikelets (Davie et al., 2008). Thus, we performed whole-cell recordings of CF EPSCs in the presence of low concentrations of the α-amino-3-hydroxy-5-methyl-4-isoxazolepropionic acid receptor (AMPAR) antagonist, NBQX (0.5 µM), to facilitate voltage clamp control (as in Dittman and Regehr, 1998; Liu and Friel, 2008; Rudolph et al., 2011).

Peak EPSC amplitudes were 37% larger in α2δ-2 KO mice (Figures 2A and 2B; WT = 703 ± 63 pA, n = 17; KO = 961 ± 86 pA, n = 19, p = 0.03; unpaired Student’s t-test). WT and KO EPSC decays were well-fit by a single exponential curve with KO EPSCs exhibiting faster decay kinetics (Figures 2C and 2D; WT τ_decay_ = 20.6 ± 1.1 ms, n = 17; KO τ_decay_ = 13.2 ± 0.8 ms, n = 20, p < 0.0001; unpaired Student’s t-test). Despite these two alterations, the EPSC in the KO had an equivalent charge transfer to that of WT EPSCs (Figures 2E), and 20-80% risetimes were similar in WT and KO PCs (Figures 2F and 2G).

**Figure 2.**
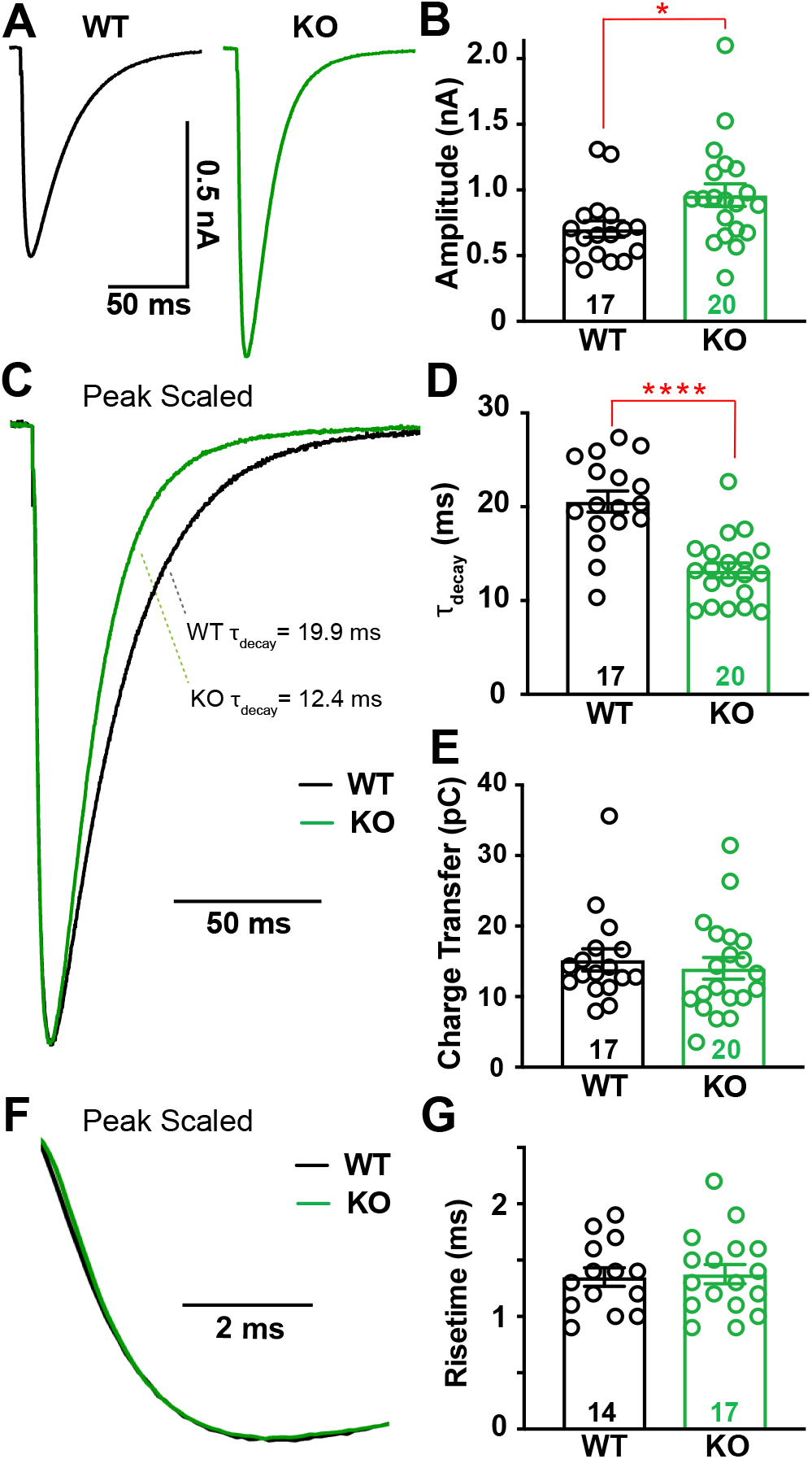
CF-evoked EPSCs are larger and faster in Cacna2d2 KO mice, but total charge transfer is conserved. (*A*) Representative CF-evoked EPSCs. *Left* WT average (black); *Right* KO average (green). (*B*) Average peak CF EPSC amplitude; *p < 0.05. (*C*) Peak scaled EPSCs, demonstrating the relative decay time constants for these example traces (τ_decay_) based on single exponential fits; WT (black) and KO (green). (*D*) τ_decay_ (ms) for CF EPSCs in WT vs. KO PCs; **** p< 0.0001. (*E*) Average charge transfer within the first 100 ms of EPSC; p = 0.58 (NS). (*F*) Peak scaled EPSCs, expanded to display risetime kinetics; WT (black) and KO (green). (*G*) Average CF EPSC 20-80% risetime (ms); p = 0.83 (NS). Data shown ± SEM, n = cells; unpaired Student’s t-test.

### Proximal redistribution of CF synapses in the α2δ-2 knockout partially contributes to larger CF-evoked EPSCs

To examine the larger EPSC in KO mice, we isolated quantal events at CF synapses by desynchronizing CF evoked release. We replaced extracellular calcium with strontium (Rudolph et al., 2011; Zhang et al., 2015), and measured the amplitudes of CF-derived asynchronous quantal release events (aEPSCs). The average aEPSC was 24% larger in KO compared to WT PCs (Figures 3A-C; WT = 25.6 ± 1.0 pA, n = 8; KO = 31.8 ± 1.6 pA, n = 8, p = 0.004; unpaired Student’s t-test), indicating that part, but not all, of the increased CF-evoked EPSC amplitude could be accounted for by a larger unitary response.

**Figure 3.**
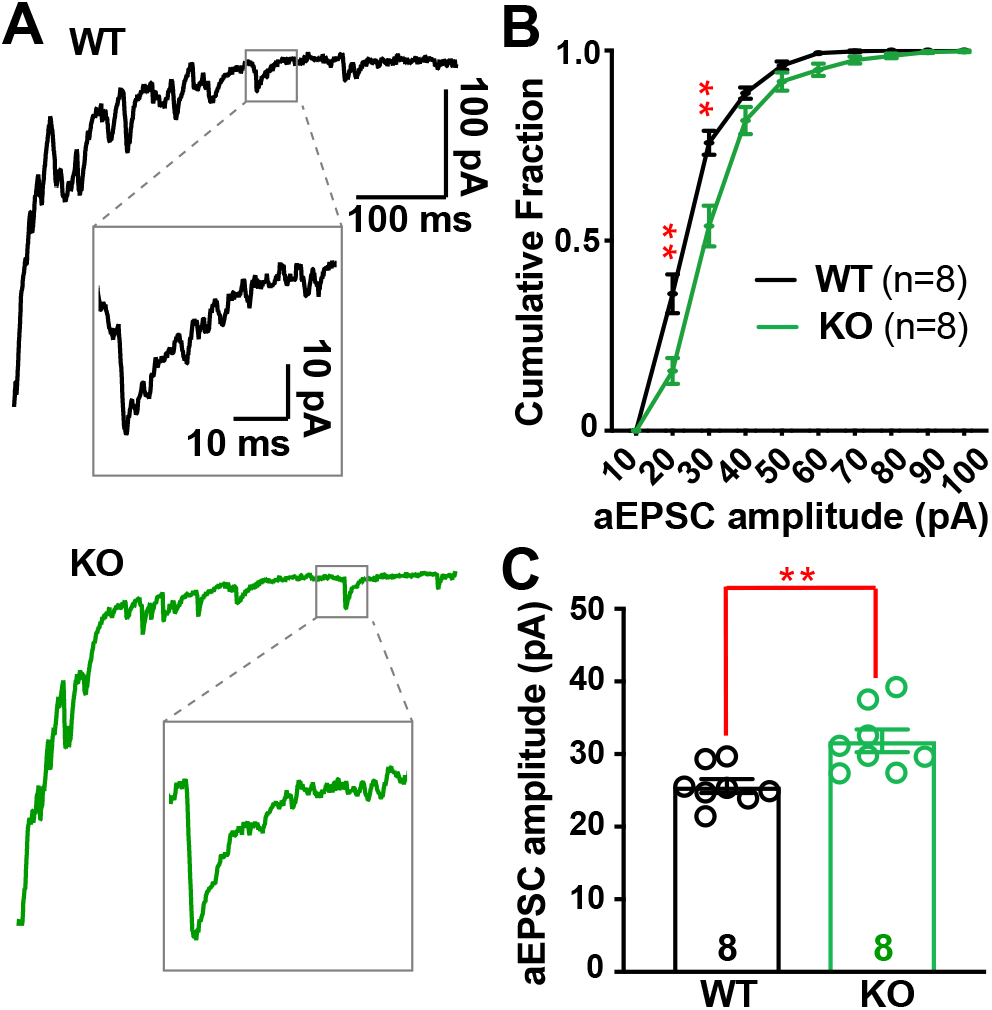
Desynchronized CF-evoked vesicle release reveals larger quantal responses in Cacna2d2 KO. (*A*) Representative CF-evoked EPSCs in the presence of 1.3 mM Sr^2+^; *Top* WT EPSC (black) and example asynchronous EPSC (aEPSC; inset); *Bottom* KO EPSC (green) and aEPSC (inset). (*B*) Cumulative aESPC amplitude distribution graphed in 10 pA bins; WT (black) and KO (green); **p < 0.01 for 20 and 30 pA bins, all others NS (Multiple t-tests with Holm-Sidak correction for multiple comparisons). (*C*) Average aEPSC amplitudes; **p < 0.01. Data shown ± SEM, n = cells; unpaired Student’s t-test.

Therefore, we asked whether there might also be an increase in the number of CF synapses onto PCs using an immunohistochemical approach. CF terminals can be selectively identified as discrete puncta along the primary dendrites of PCs through their expression of VGLUT2 (Miyazaki et al., 2004; Zhang et al., 2015). As is apparent in Figure 4A, VGLUT2^+^ puncta were closer to PC somata in KO mice than in WTs. CF terminals were a mean distance of 40.8 ± 0.5 µm from the PC soma in WT, whereas α2δ-2 KO mice had a mean CF terminal distance of 30.6 ± 0.3 µm (Figures 4A and 4C; n = 3-5 images from 5 mice of each genotype; p < 0.0001; Kolmogorov-Smirnov test). Moreover, distal CF innervation (beyond 50 µm from the PC soma; Figure 4B) accounted for 23% of puncta in WT, but only 6% in KO. This shift in CF terminal distribution in KO animals was not associated with a change in VGLUT2^+^ puncta size (Figure 4D) or overall number of puncta within the molecular layer (Figure 4E; Mean number of puncta normalized to length of PCL: WT = 1.91 ± 0.21puncta/µm_PCL_; KO = 1.81 ± 0.14 puncta/µm_PCL_, n = 3-5 images from 5 mice of each genotype; p = 0.7; unpaired Student’s t-test). There was no change in the molecular layer width, and the density of PCs was unchanged (Mean molecular layer width; WT = 110 ± 3.8 µm; KO = 106 ± 4.8 µm, p = 0.43; Mean PC density; WT = 0.62 ± 0.08 cells/µm_PCL_; KO = 0.65 ± 0.06 cells/µm_PCL_, p = 0.72; n = 8 images from 3 mice of each genotype; unpaired Student’s t-test).

**Figure 4.**
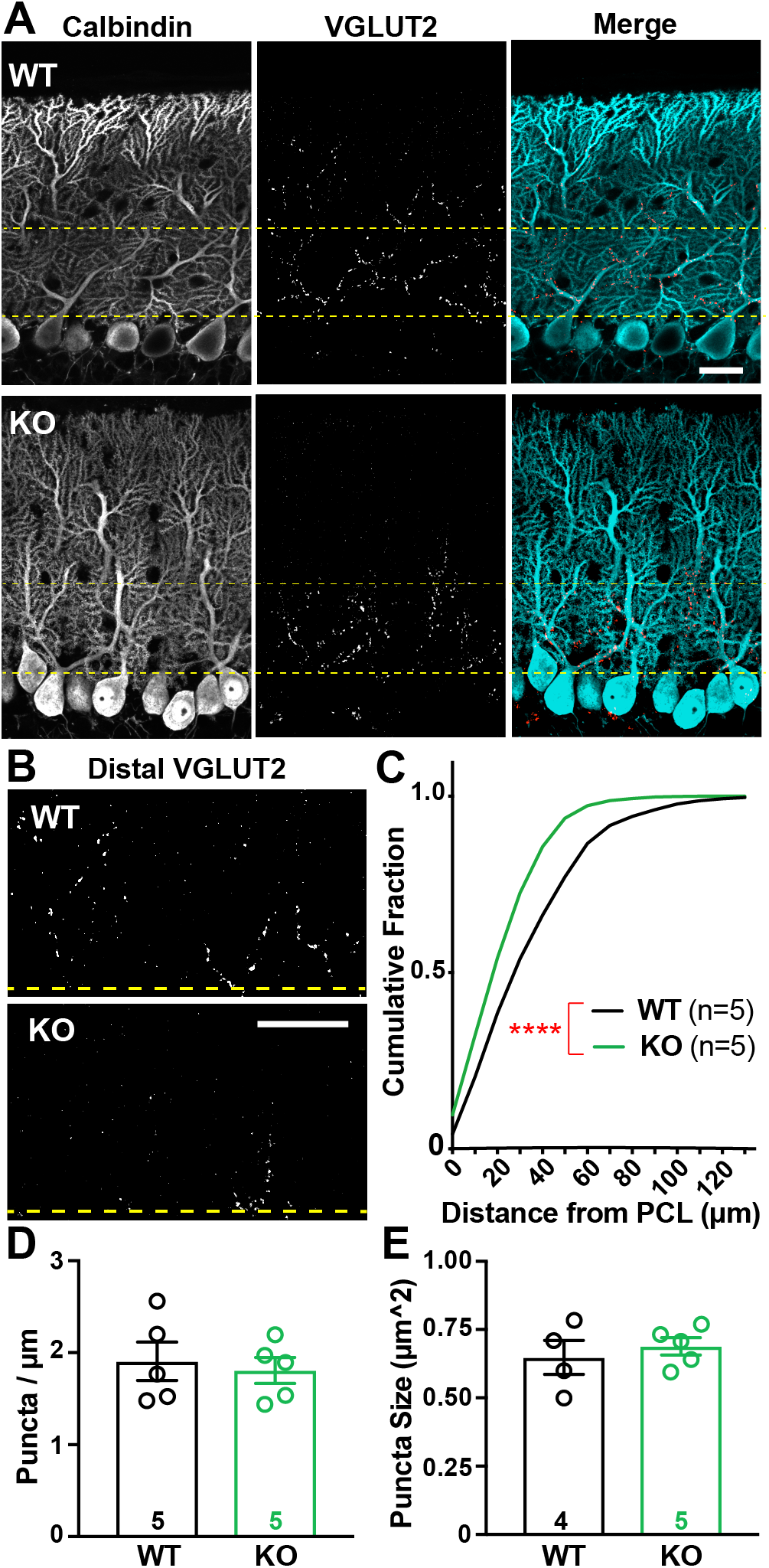
CF terminal distribution, but not number, is altered in Cacna2d2 KO cerebellum. (*A*) Representative images from p21 WT (above) and KO (below) tissue, depicting the Purkinje cell layer (PCL). Calbindin (left/blue in merge) marks PCs, while VGLUT2 immunoreactivity (middle/red in merge) marks CF terminals. Yellow lines demarcate the 50 µm most proximal to PC somata and is the region most highly innervated by climbing fibers. Scale bar = 20 µm. (*B*) VGLUT2-immunoreactive CF terminals in the outer molecular layer, cropped at the distal yellow line (50 µm), illustrate differences in CF innervation of distal PC dendrites in WT (top) and KO (below) PCs. Scale bar = 20 µm. (*C*) Cumulative distribution of VGLUT2^+^ puncta relative to PC somas in WT (black, n = 5 animals) and KO (green, n = 5 animals); ****p < 0.0001 (Kolmogorov-Smirnov test). (*D*) Average VGLUT2^+^ punctum size was not significantly different between WT and KO terminals; p = 0.55 (NS). (*E*) Average VGLUT2^+^ puncta density per length of PCL (puncta/ µm_PCL_) was not significantly different between WT and KO; p = 0.72 (NS). Unless otherwise stated, data shown ± SEM, n = animal; unpaired Student’s t-test.

The more proximal location of CF inputs in the KO could contribute to both the increased EPSC amplitude and decay rate, due to decreased dendritic filtering (Roth and Häusser, 2001). To ask whether changes in CF synapse localization alone were sufficient to account for the altered EPSC amplitude and kinetics, we modified the Roth and Häusser (2001) computational model of dendritic integration of CF inputs onto PCs to match our data. This model, based on morphological reconstructions of single PCs and empirical measurements of dendritic filtering (Roth and Häusser, 2001), allowed us to simulate how redistribution of CF inputs would affect the ensemble CF EPSC. Using this model (Figures 5A; For KO, “Model CF_70% Control_” simulates the empirically observed VGLUT2^+^ distribution from Figure 4C), a shift from the WT to the KO distribution of CF inputs produced a 16% increase in simulated EPSC amplitude and a 15% decrease in the decay time constant (Figures 5B). Thus, the proximal distribution of CF inputs in the KO augment the quantal response (Figure 5C), to account for the increase in evoked CF EPSC amplitude.

**Figure 5.**
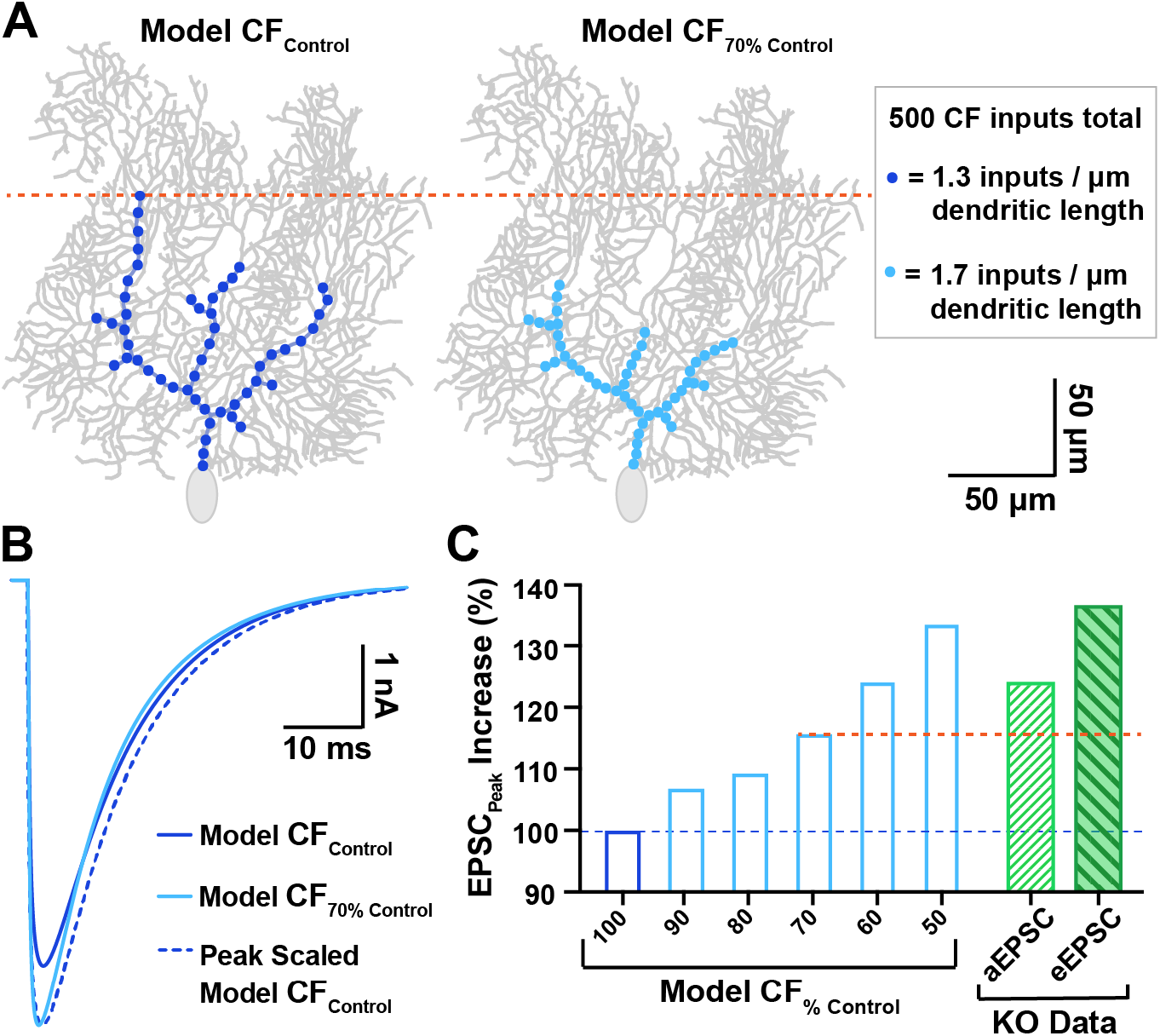
Computational PC model simulates the impact of proximally shifted CF inputs on EPSC waveform. (*A*) *Left* Model CF input distribution similar to control PCs (dark blue; “Model CF_Control_”) vs. a similar PC with CF inputs shifted 30% more proximal (*right,* light blue, “Model CF_70% Control_”), which matches the degree of proximal shift in WT vs. KO innervation, respectively. All models conserved the total number of CF quantal inputs (500 inputs with 1 nS conductance), though input density was adjusted to accommodate the shortened region of CF innervation (see inset). (*B*) Overlay of EPSC output waveforms from Model CF_Control_ simulations (dark blue; 4.7 nA), Model CF_70% Control_ (light blue; 5.4 nA), and peak scaled Model CF_Control_ to compare decay kinetics. For tau of decay; Model CF_Control_ τ_decay_ = 12.0 ms; Model CF_70% Control_ τ_decay_ = 10.2 ms). (*C*) Predicted increase in EPSC peak amplitude for various degrees of proximally shifted Model CFs (light blue bars, restricted to a zone 100-50% the width of control CFs, all including 500 quantal inputs) compared to Model CF_Control_ (dark blue bar). For comparison, the empirically determined increase in quantal EPSC (aEPSC; hatched green bar) and evoked EPSC (eEPSC; filled-hatched dark green bar) amplitudes in KO PCs are also displayed (derived from Figures 3 and 2, respectively). The orange dotted line demarcates the predicted EPSC increase from the model based on the observed shift in CF location.

### Increased multivesicular release from α2δ-2 knockout CFs

CF synapses exhibit multivesicular release, increasing the synaptic glutamate concentration (Wadiche and Jahr, 2001; Rudolph et al., 2011). To determine whether KO mice displayed altered multivesicular release, we used the low affinity, competitive AMPAR antagonist, kynurenic acid (KYN), to assay synaptic glutamate concentrations. Because KYN binds and unbinds AMPARs throughout the duration of the CF-evoked glutamate transient, KYN inhibition of the AMPAR-mediated current is inversely proportional to the concentration of glutamate present at postsynaptic receptors (Wadiche and Jahr, 2001). KYN (1µM) inhibited WT EPSC peak amplitudes by 65% (Figures 6A and 6B; EPSC_Peak_ amplitude control vs. KYN; WT_Control_ = 1.77 ± 0.41 nA vs. WT_KYN_ = 0.62 ± 0.16 nA, n = 6, p = 0.007; paired Student’s t-test), whereas KO EPSCs were inhibited by only 40% (EPSC_Peak_ amplitude control vs. KYN; KO_Control_ = 2.63 ± 0.31 nA vs. KO_KYN_ = 1.57 ± 0.21 nA, n = 8, p = 0.0009; paired Student’s t-test; relative change in WT vs. KO, p = 0.001; unpaired Student’s t-test), demonstrating enhanced multivesicular release from KO CFs.

**Figure 6.**
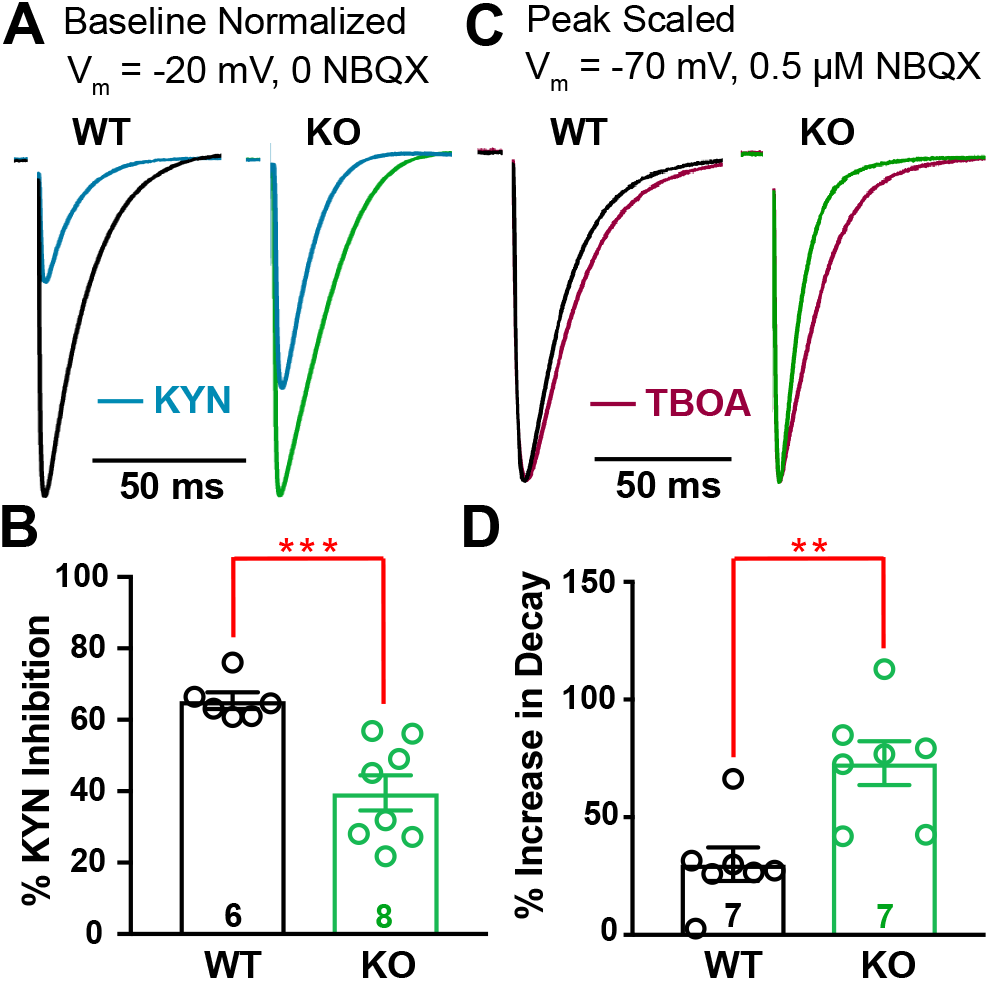
Cacna2d2 KO has increased glutamate release and clearance at CF-PC synapses. (*A*) Representative CF EPSCs recorded at V_m_ = −20 mV in the absence of NBQX for WT and KO PCs (relative scales; WT = black; KO = green). For each, traces after exposure to 1 mM kynurenic acid (KYN; blue) are shown normalized to baseline EPSC amplitudes. (*B*) Percent inhibition of ESPC peak amplitude by KYN; ***p < 0.001. (*C*) Representative normalized CF-evoked EPSCs recorded at V_m_ = −70 mV in the presence of 0.5 µM NBQX; *Left* WT average (black); *Right* KO average (green). Overlay average peak-scaled traces after exposure to 50 µM DL-TBOA (TBOA; magenta). (*D*) Average increase in EPSC decay by TBOA; **p < 0.01. Data shown ± SEM, n = cells; unpaired Student’s t-test.

For these experiments, PCs were held at −20 mV to maintain voltage clamp of the CF-evoked EPSC (Harrison and Jahr, 2003; Rudolph et al., 2011) and NBQX was omitted because co-application of NBQX and KYN facilitates AMPAR-mediated responses (Prescott et al., 2006). Interestingly, at this holding potential, KO EPSCs exhibited slower decay kinetics (For τ_decay_ at baseline V_m_ = −20 mV; WT_-20mV_ = 11.6 ± 1.12 ms, n = 6; KO_-20mV_ = 18.5 ± 1.32 ms, n = 8, p = 0.002; unpaired Student’s t-test). Although this is consistent with a larger glutamate transient due to multivesicular release (Paukert et al., 2010), it is in apparent odds with the faster decay kinetics in KO PCs at more hyperpolarized potentials. One explanation for this discrepancy in decay kinetics could be the voltage-dependence of glutamate re-uptake by PCs. PCs and surrounding Bergmann glia express high levels of glutamate transporters to manage spillover and glutamate clearance, and activity of these transporters shapes the CF-evoked EPSC waveform (Paukert et al., 2010). However, PCs play a dominant role in synaptic glutamate clearance (Auger and Attwell, 2000), and glutamate transporters have decreased efficiency at depolarized voltages (Bergles et al., 1997). In this scenario, the contributions of PC and Bergmann glia glutamate transporters together result in rapid EPSC decay in the KO at hyperpolarized potentials, but prolonged decay expected from multivesicular release dominates the EPSC waveform when KO PCs are held at depolarized potentials.

### Faster EPSC decay in α2δ-2 knockout due to enhanced glutamate clearance

We further investigated the role of glutamate transporters in shaping the CF EPSC waveform in WT and KO mice while holding PCs at −70 mV in the presence of low NBQX (0.5 µM), as in our prior voltage clamp experiments (Figure 2). Block of glutamate transporters with the non-selective transport re-uptake inhibitor, DL-TBOA (50 µM), increased WT decay constants by ∼30% (similar to Rudolph et al., 2011) (Figure 6C; WT τ_decay_ control vs. TBOA; WT_Control_ = 13.6 ± 0.9 ms vs. WT_TBOA_ = 17.4 ± 0.9 ms, n = 7, p = 0.001; paired Student’s t-test). In contrast, TBOA increased decay constants of KO EPSCs by 73% (KO τ_decay_ control vs. TBOA; KO_Control_ = 11.1 ± 1.1 ms vs. KO_TBOA_ = 19.1 ± 2.2 ms, n = 7, p = 0.001; paired Student’s t-test). After TBOA exposure, WT and KO had similar decay constants (τ_decay_ WT vs. KO in TBOA; p = 0.48; unpaired Student’s t-test). However, the relative effect of TBOA on decay was greater in KO EPSCs (Figure 6D; % increase in τ_decay_ WT vs. KO, p = 0.003; unpaired Student’s t-test) consistent with enhanced glutamate clearance by surrounding glutamate transporters.

### Repetitive stimulation of CFs suggests lower release probability in the α2δ-2 knockout, though cumulative vesicle release is greater

The enhanced multivesicular release in the α2δ-2 KO (Figure 6A-B) suggests a presynaptic contribution to the CF EPSC phenotype. CF synapses have high initial probability of release (P_R_), which is associated with paired-pulse depression at short interstimulus intervals (Hashimoto and Kano, 1998). However, CF-evoked EPSCs from KO mice showed a consistent increase in the paired-pulse ratio when compared with WT CFs (Figures 7A and 7B; Paired-pulse ratio: WT = 0.41 ± 0.03, n = 17; KO = 0.51 ± 0.02, n = 18, p = 0.01; unpaired Student’s t-test), suggesting CFs have a lower P_R_ when α2δ-2 is deleted. Therefore, we hypothesized that KO CFs might have a substantially greater readily releasable pool of vesicles to exhibit increased multivesicular release while also having a lower P_R_.

**Figure 7.**
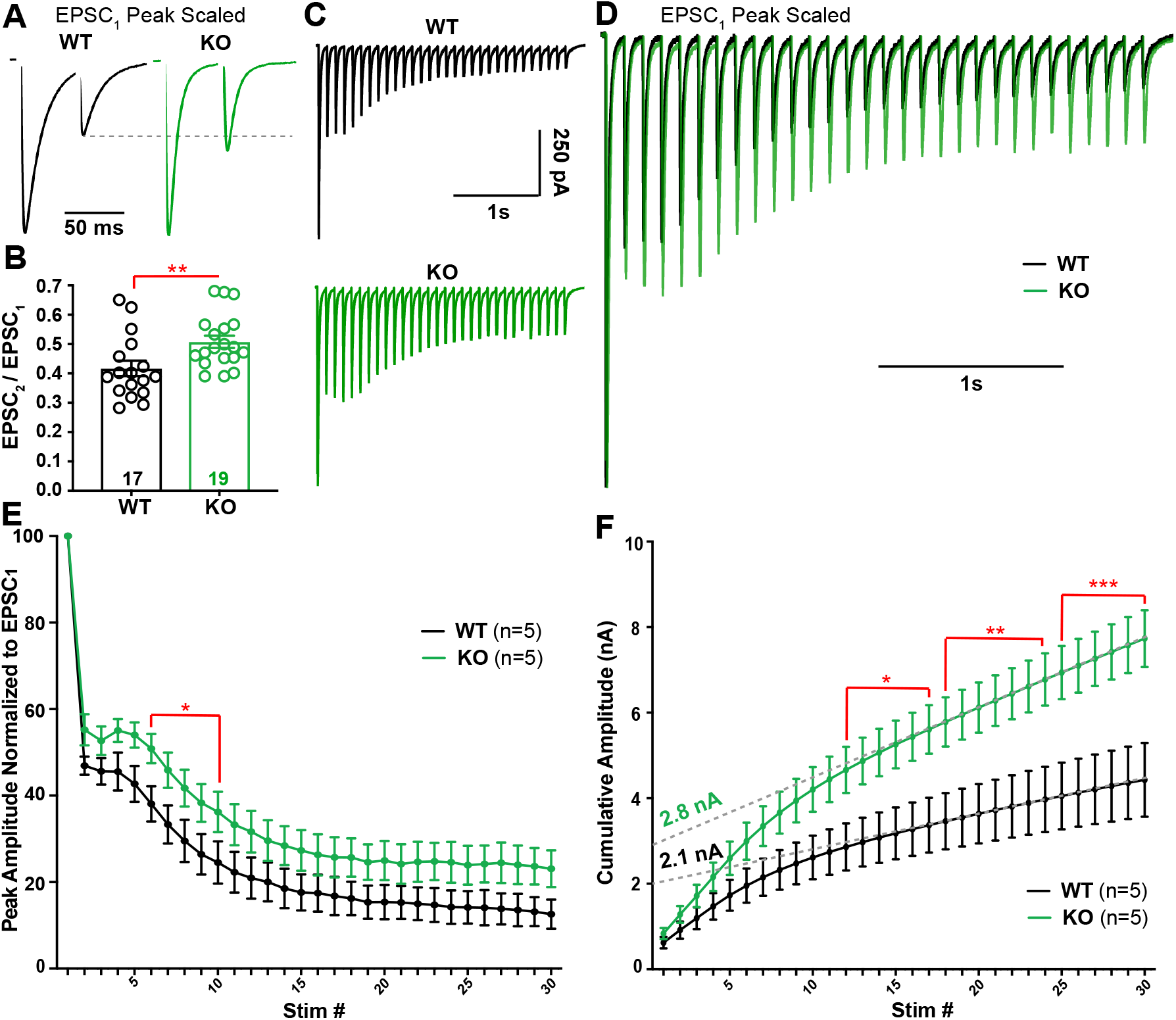
Repetitive stimulation of CF synapses reveals a lower probability of release and greater cumulative release in Cacna2d2 KO. *A*) Representative traces from WT (black) and KO (green) PCs during 50 ms paired-pulse stimulation. Traces are scaled to the first EPSC (EPSC_1_). Dotted gray line shows paired-pulse depression of the second EPSC (EPSC_2_) in WT compared to KO. (*B*) Average paired-pulse ratio (EPSC_2_ /EPSC_1_); ** p < 0.01. (*C*) Representative traces in response to 10 Hz stimulation; WT (black); KO (green). (*D*) Traces from (*C*) peak scaled to EPSC_1_ and overlaid, illustrating different relative steady-state EPSC amplitudes during latter portions of the train. (*E*) Summary data of EPSC amplitudes normalized to EPSC_1_ during 10 Hz stimulation in WT (black) and KO (green). *p < 0.05 (Multiple t-tests with Holm-Sidak correction). (*F*) EPSC amplitudes during 10 Hz stimulation plotted as cumulative amplitude from WT (black) and KO (green). For comparison of cumulative amplitude between WT and KO at various stimulation numbers (stim #); * p < 0.05, ** p < 0.01, *** p < 0.001 (Multiple t-tests with Holm-Sidak correction). Dotted grey lines illustrate a linear fit to cumulative amplitude between stim # 20-30 from WT and KO trains. Data shown ± SEM, n = cells; unpaired Student’s t-test unless otherwise indicated.

To estimate the relative size of the readily releasable pool in WT and KO, we stimulated CFs with a 10 Hz train. WT and KO CF synapses had strikingly different responses during repetitive stimulation. Both genotypes exhibited a delayed facilitation followed by depression, which may indicate multiple pools of vesicles with differing release probabilities (Lu and Trussell, 2016). However, KO EPSCs were larger at every stimulus throughout the train (Figures 7C-E), providing further evidence of enhanced vesicle release compared to WT. A linear fit to the last 10 responses of a cumulative amplitude plot produced a steeper slope (Linear regression using 95% Confidence Intervals; WT_slope_ = 78.7 ± 0.6, n = 5; KO_slope_ = 172.2 ± 0.5, n = 5; p < 0.0001; unpaired Student’s t-test) and a larger y-intercept in KO PCs when compared to WT (Figure 7F). This analysis provided an estimate of the readily releasable pool (Schneggenburger et al., 1999), which was 29% larger in the KO compared to WT (WT y-intercept = 2.08 ± 0.02 nA, n = 5; KO y-intercept = 2.77 ± 0.01 nA, n = 5; p < 0.0001; unpaired Student’s t-test). Thus, both a single stimulus (Figure 6A-B) and repetitive stimulation of CFs produced increased vesicle release in the α2δ-2 KO, suggesting either an enhancement of a low P_R_ pool of vesicles (Lu and Trussell, 2016) and/or more discrete release sites with reduced P_R_ compared to WT.

### CF terminals in α2δ-2 knockout have more release sites

Because we did not see an increased density of VGLUT2^+^ puncta by light microscopy, we hypothesized CF terminals in the KO had either a greater number of release sites or vesicles. CF terminals can be identified with electron microscopy (EM) by their high density of round synaptic vesicles (SVs) and contacts onto PC “thorns” along primary PC dendritic shafts (Palay and Chan-Palay, 1974; Miyazaki et al., 2004). While blinded to genotype, we imaged and quantified CF terminals from WT and KO mice (10-15 images/animal; see Materials and Methods). In agreement with VGLUT2^+^ puncta size determined by confocal microscopy (Figure 4D), CF terminal cross-sectional area was no different between genotypes when sampled using EM (Figures 8A and 8B). WT and KO CF terminals also had an equivalent density of synaptic vesicles (SVs) (Figure 8C) and no change in number of SVs within 100 nm of the synaptic contact, a proxy for the readily releasable pool (WT = 18.5 ± 2.4 SV/µm_Contact_, n = 5 mice; KO = 20.4 ± 2.7 SV/µm_Contact_, n = 6 mice, p = 0.6; unpaired Student’s t-test).

**Figure 8.**
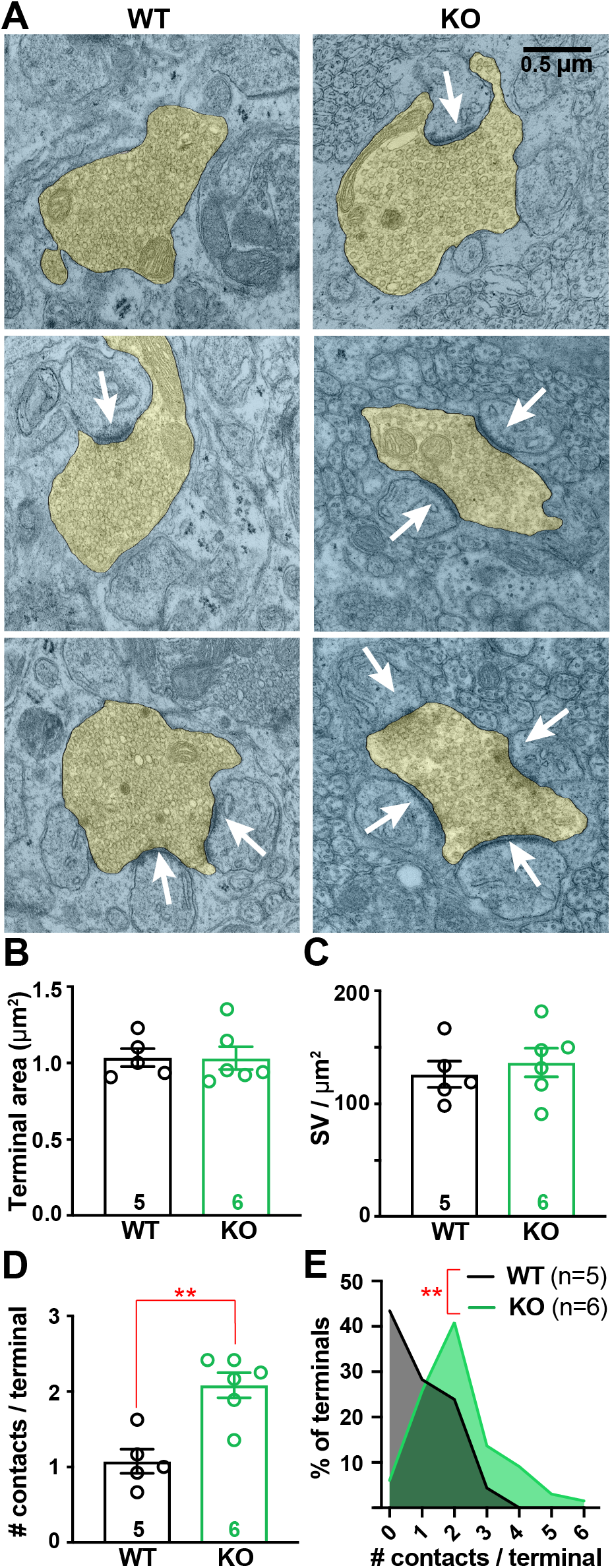
CF terminals have increased numbers of synaptic contacts in Cacna2d2 KO animals. (*A*) Representative transmission electron micrographs of CF terminals (pseudocolored yellow) from p21 WT (left) and KO (right) animals. White arrows indicate postsynaptic densities used to quantify synaptic contacts/terminal. Scale bar = 0.5 µm. (*B*) Average CF terminal area (µm^2^); p = 0.55 (NS). (*C*) Synaptic vesicle (SV) density (SV/ µm^2^) was not different between WT and KO animals; p = 0.57 (NS). (*D*) Average number of contacts per CF terminal (# contacts/terminal) was increased in KO animals. ** p <0.01 (Mann-Whitney test). (*E*) Histogram of all CFs analyzed from WT (black) and KO (green) cerebelli, displaying the number of contacts per sampled CF terminal normalized to total number of CF terminals; p < 0.01 (Kolmogorov-Smirnov test). Data shown ± SEM, n = animals using 15-20 images/animal; unpaired Student’s t-test unless otherwise stated.

However, the number of synaptic contacts made by CF terminals in α2δ-2 KOs was nearly twice that of WT (Figure 8A **and** 8D; WT = 1.08 ± 0.16 contacts/terminal, n = 5 mice; KO = 2.08 ± 0.17 contacts/terminal, n = 6 mice, p = 0.009; Mann-Whitney test). Moreover, the distribution of the number of contacts made by CF terminals was right-shifted in KO (Figure 8E; p < 0.002; Kolmogorov-Smirnov test). Although quantifying synaptic contacts per terminal using single EM sections likely underestimates the true number of contacts per terminal, the observed doubling of discrete contacts made by KO CFs is consistent with increased multivesicular release in the α2δ-2 KO.

### Differential *Cacna2d* isoform expression in the inferior olive and Purkinje cell layer

By *in situ* hybridization, there is robust and exclusive expression of the *Cacna2d2* isoform in PCs, and relatively low expression in the inferior olive (IO), which gives rise to climbing fibers (Cole et al., 2005; Lein et al., 2007). Given the presynaptic morphological and physiological changes we observed at *Cacna2d2* KO CF-PC synapses, we re-examined expression of α2δ isoforms in the Purkinje cell layer (PCL) and IO using quantitative PCR of fresh-frozen tissue obtained by laser capture microdissection (Figure 9A). In agreement with prior observations (Cole et al., 2005; Lein et al., 2007), PCL tissue from wildtype mice expressed the α2δ-2 isoform 3.85 ± 0.96-fold higher than presynaptic IO samples (Figure 9B; *Cacna2d2* ΔCt _PCL_ = 1.62 ± 0.17; ΔCt _IO_ = 3.37 ± 0.38, n = 4, p = 0.03; paired Student’s t-test). PCL tissue also contained low expression of α2δ-1 and high levels of α2δ-3 transcript due the inclusion of other cell types in the sample, such as Bergmann glia and molecular layer interneurons, which can be appreciated by expression patterns apparent from *in situ* hybridization (Cole et al., 2005; Lein et al., 2007). Consistent with previous reports, α2δ-4 was undetectable in both PCL and IO samples. Importantly, analysis of *Cacna2d* transcripts from IO samples revealed that α2δ-1 is the predominant isoform (Figure 9C), and is 7.0 ± 2.4-fold more abundant than α2δ-2 in IO samples (*Cacna2d1* ΔCt _IO_ = 0.94 ± 0.4; *Cacna2d2* ΔCt _IO_ = 3.37 ± 0.38, n = 4, p = 0.04; paired Student’s t-test). These results suggest that the locus of action of α2δ-2 at CF-PC synapses is predominately postsynaptic.

**Figure 9.**
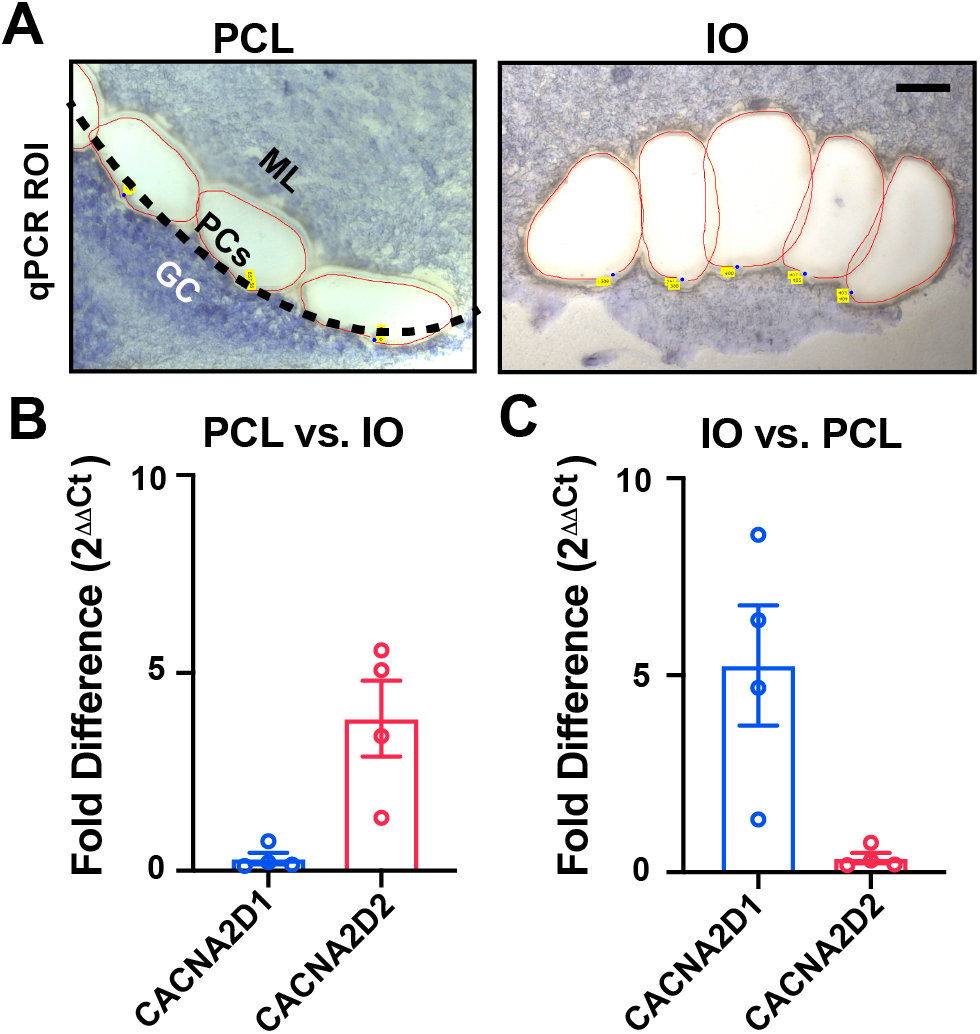
Relative expression of Cacna2d transcripts by qPCR from Purkinje cell layer and inferior olive tissues. (*A*) Example Purkinje cell layer (PCL) and inferior olive (IO) regions of interest isolated by laser capture microdissection from fresh-frozen WT tissue. *Left, PCL region of interest;* black-dotted line indicates the monolayer of PCs that have been dissected along with regions of the inner molecular layer (ML; granule cells, GC). *Right, IO region of interest;* one hemisphere from a coronal section of ventral brainstem, scale = 100 µm. (*B,C*) Fold difference (2^-ΔΔCt^) expression of *Cacna2d* isoforms by quantitative PCR comparing (*B*) PCL vs. IO, and (*C*) IO vs. PCL samples. Data shown ± SEM, n = 4 animals.

## Discussion

In mice lacking α2δ-2, profound deficiencies in CF-induced complex spike (CpS) generation were associated with altered underlying CF-evoked EPSC amplitude and kinetics. CpSs are initiated by a AMPAR-mediated depolarization that drives an initial sodium spike followed by multiple axonally generated ‘spikelets’ (Davie et al., 2008). Furthermore, dynamic clamp experiments suggested that increased CF EPSC amplitudes would result in depolarization block of spikelet generation (Davie et al., 2008). Thus, we hypothesize that increased glutamatergic transmission and accelerated EPSC kinetics (Rudolph et al., 2011) shape the CpS in the α2δ-2 KO (Figure 10).

**Figure 10.**
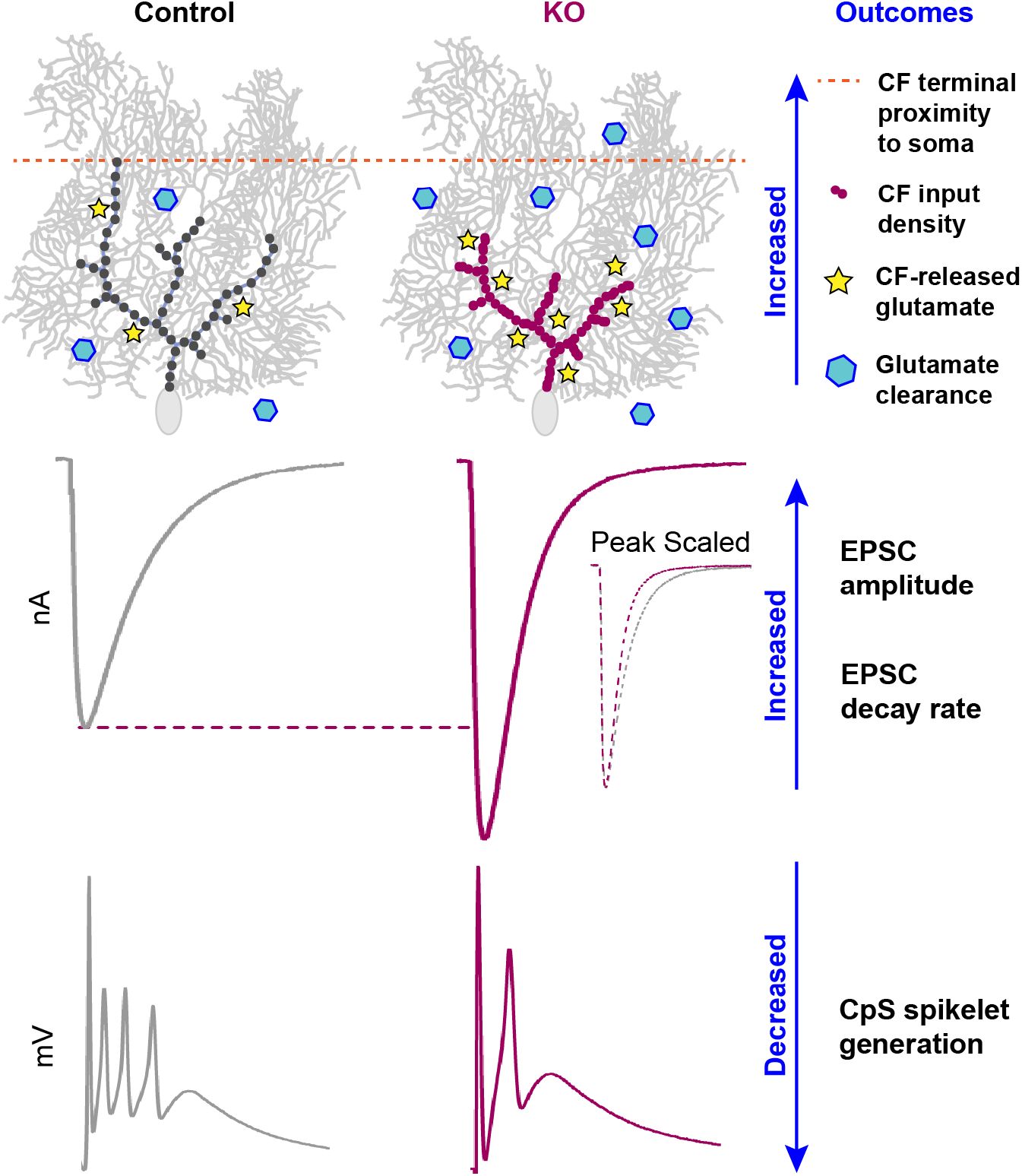
Summary of CF-PC phenotypes in Cacna2d2 KO mice. Proximal distribution of CF inputs onto KO Purkinje cells enhanced postsynaptic quantal responses to CF glutamate release, and the increased number of synaptic release sites increased total glutamate concentration. Together, this resulted in a 140% EPSC amplitude in the KO compared to control. Counteracting effects included a lower CF P_R_ and enhanced glutamate clearance, which doubled the EPSC decay rate. Ultimately, larger synaptic conductances in the KO likely contribute to depolarization block of CF-evoked spikelet generation.

Each CF terminal in the KO contained more discrete synaptic contacts, leading to increased multivesicular release and consequently larger EPSCs, despite an apparent reduction in the probability of release. Together, these findings seem to contradict observations in which synapse formation is positively regulated by α2δ expression (Li et al., 2004; Eroglu et al., 2009; Chen et al., 2018; Risher et al., 2018), reflecting a more complex and nuanced role of α2δ proteins in the α2δ-2 KO phenotype.

### α2δ-2 proteins as auxiliary Ca^2+^ channel subunits

α2δ proteins were first identified as auxiliary subunits of voltage-dependent Ca^2+^ channels (VDCCs) that facilitate surface trafficking of VDCCs in heterologous expression systems (Canti et al., 2005; Cassidy et al., 2014). α2δ-1 knockdown reduces presynaptic vesicle release in cultured hippocampal neurons (Hoppa et al., 2012), and capacitive measurements from inner hair cells of the spontaneous *Cacna2d2* mutant, *du*/*du*, show reduced exocytosis (Fell et al., 2016). We also observed a lowered release probability (P_R_) at the CF-PC synapse, as determined by paired-pulse ratio, which is consistent with these prior observations and could relate to altered VDCC localization or abundance in CFs. Given that α2δ-1 is the predominant isoform expressed by inferior olivary cells (Cole et al., 2005; Lein et al., 2007), if the phenotype we observed is mediated by presynaptic loss of α2δ-2, it suggests isoform-specific functions that cannot be compensated by α2δ-1. Furthermore, this raises the possibility that different α2δ isoforms in individual neurons may have distinct functional roles.

In addition to their abundant α2δ-2 expression, PCs highly express the VDCC, Cav2.1 (Barclay et al., 2001a), which localizes to PC somata and primary dendrites in scattered and clustered patterns (Indriati et al., 2013). Dissociated PCs from the spontaneous mutant, *du^2j^/du^2j^*, have ∼30% reduced somatic calcium currents (Barclay et al., 2001a). Similarly, CF vesicle release is mostly (70-90%) regulated by Cav2.1 (Regehr and Mintz, 1994). Because CF-PC transmission and development has been extensively characterized in Cav2.1 mutant mice (Matsushita et al., 2002; Miyazaki et al., 2004; Hashimoto et al., 2011), we looked to this literature for clues as to whether our observed phenotypes could reflect altered pre- and/or postsynaptic VDCC function or localization.

For example, the *leaner* phenotype, in which a Cav2.1 pore mutation reduces PC calcium currents by 60%, is associated with larger, rapidly decaying CF-evoked EPSCs (Liu and Friel, 2008), like the α2δ-2 KO. However, *leaner* CFs do not exhibit changes in P_R_ (Liu and Friel, 2008). Other Cav2.1 pore mutants show reductions in PC calcium current analogous to α2δ-2 mutants, but display heterogeneity in EPSC phenotypes. Both *rolling Nagoya* and *tottering* mutants have 40% reductions in calcium currents, yet *rolling Nagoya* exhibits larger, slowly decaying EPSCs, whereas *tottering* EPSCs are unchanged (Matsushita et al., 2002). PC-specific Cav2.1 KO show proximal innervation by CFs (Miyazaki et al., 2012), reminiscent of the CF redistribution we observed in *Cacna2d2* KOs. This striking similarity provides evidence for postsynaptic control of CF development. Surprisingly however, global and PC-specific Cav2.1 KOs exhibit normal CF EPSC amplitude and decay kinetics (Miyazaki et al., 2004; Hashimoto et al., 2011). Thus, although some similarities exist between the *Cacna2d2* KO and certain VDCC mutant mice, the roles of α2δ-2 at the CF-PC synapse likely involve other effector mechanisms. α2δ-2 loss may be more nuanced than altered VDCC abundance, but perhaps result from differences in VDCC localization, clustering or association with other molecules (including other VDCC subtypes).

Altered presynaptic function could result from PC-driven morphological changes in the *Cacna2d2* KO, independent of altered presynaptic VDCC trafficking. For example, alterations in presynaptic morphology alone, rather than altered presynaptic gene expression, can influence measures of apparent P_R_ and the readily releasable pool size by repetitive stimulation (Fekete et al., 2019). It is also possible that P_R_ at individual release sites is unchanged, but an increased number of release sites per terminal could allow for accumulation of [Ca^2+^]_i_ during repetitive stimulation via inter-site crosstalk. In this scenario, repetitive stimuli recruit vesicles from a low P_R_ pool, sustaining release during subsequent stimuli. Currently, there is no definitive way to distinguish between a heterogeneous population of P_R_ at discrete release sites, or whether the vesicle recruitment from low vs. high P_R_ pools differs (Kaeser and Regehr, 2017). The possibility of multiple vesicle pools in the CF, as seen by the bimodal responses in Figure 7C, complicates our ability to derive readily releasable pool size and P_R_ at this synapse (Neher, 2015; Lu and Trussell, 2016). Given the various possible effector mechanisms for α2δ-2 that could contribute to the phenotype in mutant mice, further experiments using cell-type selective genetic manipulations will be necessary to fully understand the locus of α2δ-2 action.

### α2δ-2 proteins as synaptic organizers

Recently, roles for α2δ proteins independent of VDCCs are supported by their ability to regulate synapse formation despite pharmacological block or deletion of VDCCs (Eroglu et al., 2009; Kurshan et al., 2009). α2δ-1 induces excitatory synapse formation in response to glial-secreted thrombospondin (Eroglu et al., 2009), which is dependent on postsynaptic signaling through N-methyl-D-aspartate receptors (NMDARs) (Risher et al., 2018). Though NMDARs are transiently expressed in newborn rat PCs (Rosenmund et al., 1992), they are not detected at mouse CF-PC synapses until late adulthood (Piochon et al., 2007; Renzi et al., 2007), making this interaction unlikely to contribute to the *Cacna2d2* KO phenotype.

Studies in *C. elegans* suggest trans-synaptic roles for α2δ proteins, as presynaptic α2δ-3 and binds to postsynaptic neurexin to control neuromuscular synaptic function (Tong et al., 2017). In mammals, neurexins are presynaptically expressed (Missler et al., 2003; Zhang et al., 2015) and interact trans-synaptically with postsynaptic neuroligins. Interestingly, similar to our observations in the *Cacna2d2* KO, the neuroligin triple KO mouse (Zhang et al., 2015) also has a proximally-shifted CF distribution, without a change in VGLUT2^+^ puncta size or overall density (Zhang et al., 2015). Although CF-PC synaptic function is unchanged in the neuroligin tKO (Zhang et al., 2015), this finding suggests that α2δ-2 may act in parallel with neurexin-neuroligin to coordinate features such as synaptic localization, whereas the functional components that depend on α2δ-2 involve other effector mechanisms.

The stereotyped development of the CF-PC synapse has made it an attractive model to study numerous important molecules that share mutant phenotypes, yet have quite disparate functions. For example, the CF innervation pattern is modulated by postsynaptically expressed neuroligins, TrkB, Cav2.1, GluRδ2 (exclusively expressed in PCs), cerebellins and myosin Va (reviewed in Bosman and Konnerth, 2009), revealing CF redistribution as a sensitive indicator of underlying dysfunction, and demonstrating postsynaptic control of synaptic innervation. Although *Cacna2d2* KO mice share this phenotype, additional functional alterations at the CF-PC synapse are dissimilar from other mutant mice. Thus, our data suggest that the contributions of α2δ-2 to the CF-PC synapse may comprise both VDCC-dependent and -independent mechanisms.

### Impacts of α2δ-2 loss on cerebellar function

Of the four α2δ isoforms (*Cacna2d1-4*), α2δ-2 loss has the most severe phenotype, as mice and humans with *Cacna2d2* mutations have ataxia, epilepsy and motor control deficits (Barclay et al., 2001b; Brodbeck et al., 2002; Ivanov et al., 2004; Donato et al., 2006; Pippucci et al., 2013). Because PCs provide the output from the cerebellum, some of these neurologic phenotypes likely reflect PC dysfunction. PC spike-rate and plasticity are instructed by CpSs, providing error prediction information for motor coordination, presumably graded by the number of spikelets generated (Rasmussen et al., 2013; Yang and Lisberger, 2014; Burroughs et al., 2017). As CpSs generated by *Cacna2d2* KO PCs had fewer spikelets, we predict PC information transfer is degraded, implicating direct influence of α2δ-2 loss in cerebellar dysfunction.

Particularly surprising was the variety of alterations at the *Cacna2d2* KO CF-PC synapse, including enhanced glutamate re-uptake. It is likely that some phenotypes developed as compensation for a primary derangement directly related to α2δ-2 loss. Delayed conditional deletion of α2δ may help to clarify this issue. Likewise, deletion of α2δ-2 may have differential effects on other synapses and functions, either onto PCs or other cell types. As many neurons co-express multiple α2δ isoforms, it will be important to understand the role of each isoform at individual synapses, as well as address whether postsynaptic α2δ proteins instruct presynaptic development or function (or vice versa), using cell-specific targeted rescue or deletion. Overall, our results underscore the critical roles of α2δ-2 in both proper organization and transmission at the CF-PC synapse.

## Acknowledgments

This material was supported in part by the Department of Veterans Affairs, Veterans Health Administration, Office of Research and Development, Biomedical Laboratory Research and Development Merit Review Award I01-BX002949 (ES), a Department of Defense CDMRP Award W81XWH-18-1-0598 (ES), NIH T32NS007466 (KAB), NIH R01-NS080979 (GLW), NINDS 1R21NS102948 (IK/ES), NIH P30NS061800 (Dr. Sue Aicher), and OHSU Innovation Fund (ES) awards. The electron microscope was purchased with funds from the Murdock Charitable Trust awarded to Dr. Aicher.

We thank Dr. Stefanie Kaech-Petrie of the OHSU Advanced Light Microscopy Core for assistance with imaging; Dr. Gail Mandel and Dr. John Sinnamon for expertise and use of laser capture equipment; the OHSU Gene Profiling/RNA and DNA Services Shared Resource for support with qPCR experiments; and Dr. Sue Aicher, James Carroll, and Jo Hill for the electron microscopy expertise. We would like to acknowledge Drs. Sergey Ivanov and Lino Tessarollo for generously providing the *Cacna2d2* knockout mice; Drs. Michael Häusser and Arnd Roth for necessary files and permission to use the NEURON model; and research associates Arielle Isakharov and Ada Zhang for outstanding technical support. We also thank Dr. Christopher Vaaga and Dr. Laurence Trussell for constructive comments on the manuscript. The contents of this manuscript do not represent the views of the U.S. Department of Veterans Affairs or the United States Government.

